# Grounding olfactory perception in language: Benchmarks and models for generating natural language odor descriptions

**DOI:** 10.64898/2026.03.04.709650

**Authors:** Cyrille Mascart, Khue Tran, Khristina Samoilova, Liam Thomas Storan, Tingkai Liu, Alexei Koulakov

**Author notes:** Equal contributions (Cyrille Mascart, Khue Tran).

## Abstract

Recent advances in deep learning have enabled prediction of odorant perception from molecular structure, opening new avenues for odor classification. However, most existing models are limited to predicting percepts from fixed vocabularies and fail to capture the full richness of olfactory experience. Progress is further limited by the scarcity of large-scale olfactory datasets and the lack of standardized metrics for evaluating free-form natural-language odor descriptions. To address these challenges, we introduce Odor Description and Inference Evaluation Understudy (ODIEU), a benchmark which includes perceptual descriptions of over 10,000 molecules paired with a model-based metric for evaluating free-form odor text descriptions. The model-based metric uses Sentence-BERT (SBERT) models which are fine-tuned on olfactory descriptions to allow better evaluation of human-generated odor texts. Using the fine-tuned SBERT models, we show that free-form text odor descriptions contain additional perceptual information in their syntactic structure compared to semantic labels. We further introduce CIRANO (Chemical Information Recognition and Annotation Network for Odors), a transformer-based model that generates free-form odor descriptions directly from molecular structure, thus implementing the molecular structure-to-text (S2T) prediction. CIRANO achieves performance comparable to humans. Finally, we generate human-like descriptions from mouse olfactory bulb neural data using an invertible SBERT model, yielding neural-to-text (N2T) predictions highly aligned with human descriptions. Together, CIRANO and ODIEU establish a standardized framework for generating natural language olfactory descriptions and evaluating their alignment with human perception.

Code is available at https://github.com/KoulakovLab/ODIEU

## Introduction

Olfaction remains the final frontier of our senses. While vision and audition benefit from a well-established understanding of how stimulus properties give rise to percepts, our understanding of olfaction is far less complete [1, 2]. A major obstacle lies in our limited ability to precisely describe olfactory percepts, resulting in a scarcity of perceptual data. Semantic data, where smells are described using words or sentences, offers valuable insights [1, 3]. In this study, we argue that the syntactic structure of complete sentence descriptions contains information beyond that captured by binary semantic labels that are often used in human computational olfaction. To evaluate the quality of various forms of odor descriptions, including machine-generated ones, we developed an LLM-based benchmark that compares these representations to human-generated texts.

Several methods have been used historically to describe human responses to odorants [2]. While pairwise similarity judgments have proven effective in mice [4], they yield an overly simplistic perceptual odor space in humans [2, 5]. Most of what we know about human olfactory perception comes from semantic datasets [1, 2, 6, 7]. One such method, known as profiling, involves a trained panel of observers evaluating whether a given term from a predefined and diverse dictionary applies to a specific odor [2, 6, 8]. Each odor-descriptor pair is then quantified by the proportion of panelists who agree on the association, resulting in real-valued vectors that represent odorants in a high-dimensional semantic space. Profiling data is available for hundreds of molecules [6, 8, 9]. Non-linear embedding methods have shown that profiling data resides near a space of small (D∼6) dimensionality [1, 3]. A larger but less precise source of information comes from binary semantic datasets, in which each term (e.g., “sweet”) is marked as either applicable or not, without indicating intensity. Such datasets include Leffingwell [10], GoodScents [11], Arctander [12], and Flavornet [13], collectively covering thousands of molecules. The corresponding molecular structures, including 3D conformations and additional chemical properties, are available through publicly accessible databases, such as PubChem [14]. Large fraction of this data was combined in the Pyrfume platform [15]. Binary semantic data contains sufficient information to train deep learning models, ranging from equivariant convolutional neural networks (CNNs) [16, 17] to transformer-based architectures [18], to predict semantic labels from molecular structure, achieving high performance levels.

Odor spaces are actively studied constructs that represent olfactory sensory objects within a shared set of dimensions, analogous to the color space in vision [1, 3, 5-7, 16, 19-24]. In color perception, for example, stimuli spanning an infinitely-dimensional light spectrum can be effectively captured by just three coordinates: hue, saturation, and intensity. Similarly, it has been proposed that, despite the existence of millions of volatile compounds, the dimensionality of olfactory perceptual space is limited [1, 3]. For example, it was proposed that one of the prominent dimensions of the odor space could be described as the odorants’ pleasantness [1, 3]. Other dimensions have also been described, including the molecules’ hydrophobicity and complexity [1]. It was proposed that the dimensions of the perceptual odor space could be associated with distinct metabolic pathways generating the smells [25]. Deep learning approaches have been used to derive low-dimensional embeddings of odorant percepts [6, 16-18]. As in vision or audition, a precise definition of odor space would be a critical step toward a deeper understanding of olfactory processing. Our present study aims at refining the definition of olfactory percepts beyond simple labels by including syntactic information present in several datasets.

In recent years, several deep learning approaches have been developed to predict olfactory perception of single-molecule odorants. Equivariant convolutional neural networks (CNNs) have been first employed to predict individual dimensions of olfactory perceptual space based on 3D molecular structures [16], and, later, extended to be predictive of discrete semantic labels [17]. Graph neural networks (GNNs) have modeled molecules as graphs to infer semantic descriptors [6, 26], while transformer networks have utilized SMILES string representations for the same purpose [18]. These models achieve high-quality structure-to-percept predictions, typically reaching AUC values above 0.9. However, it remains unclear whether such performance adequately captures the complexity of olfactory perception. Here, we argue that incorporating free-form, sentence-level descriptions of odors can significantly expand the representational capacity of odor space, capturing aspects of perception that go beyond what is measurable by AUC alone.

Some of the datasets mentioned above include not only semantic labels but also free-form text descriptions of percepts evoked by molecules (Fig. 1A). These descriptions contain syntactic structure that may carry additional information relevant to predicting olfactory percepts. In this study, for the first time, we will address this additional information. Our first goal is to develop a benchmark for evaluating free-form text descriptions of odors. To achieve this, we compare descriptions of the same molecule across individuals, quantifying both inter-observer similarity and variability. We propose that for machine-generated structure-to-text (S2T) predictions to match human quality, the similarity between model output and human ground truth must be comparable to the consensus among human observers. This principle forms the basis of our benchmark, the Odor Description and Inference Evaluation Understudy (ODIEU), which offers an objective implementation of the olfactory Turing test. We use ODIEU to evaluate several simple S2T predictors which use both 3D molecular structures and SMILES strings. Using ODIEU, we argue that the syntactic structure of human descriptions carries important additional information about olfactory perception, and that conventional metrics for predictive models such as AUC do not adequately capture the richness of olfactory descriptions.

**Figure 1:**
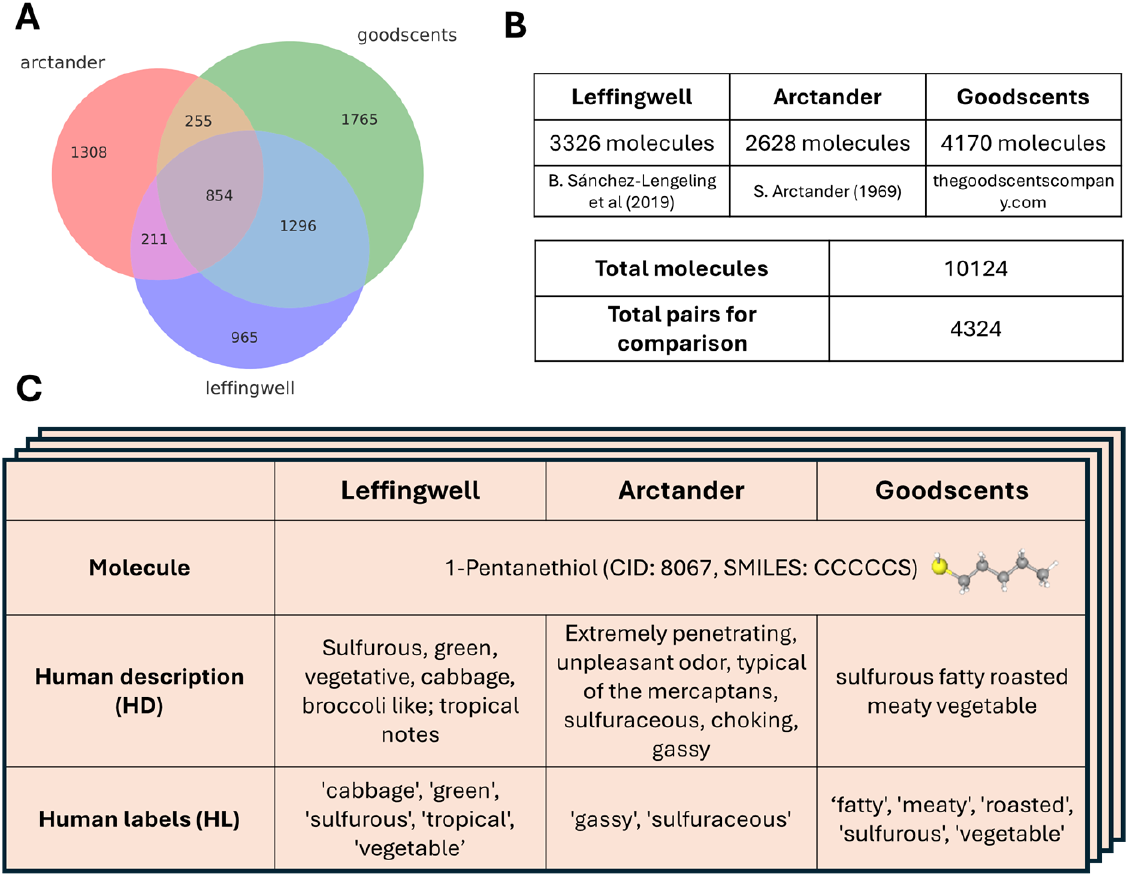
The datasets used in this study. (A) The Venn diagram of the Arctander, Leffingwell and Goodscents datasets. (B) Overview of the data. The datasets include human descriptions in the form of sentences for 10,124 molecules, including 4,324 molecules pairs of descriptions provided by two different humans for the same molecules. (C) Example of an entry for one molecule present in all three datasets (1-Pentanethiol).

After evaluating several simple S2T predictors, we trained a specialized network, called CIRANO (Chemical Information Recognition and Annotation Network for Odors), to solve the S2T problem. CIRANO is optimized to infer perceptual odor qualities from molecular structure and express them in natural language. It consists of a GPT-2 backbone with a fine-tuned encoder that projects molecular structure into the token embedding space. After evaluating CIRANO using our benchmark, we found that its performance is comparable to that of humans. Overall, using our datasets and benchmark, we developed an algorithm that targets the olfactory language grounding problem.

Finally, we designed an algorithm that uses neural data to generate odor descriptions. Because large scale neural recordings for diverse odor sets are not available in humans, we relied on neural responses to odorants collected in the mouse olfactory bulb. We used Q-vectors, neural representations recently shown to be optimal for cross-animal odor generalizations [27], to generate human natural language descriptions. As the core of this approach, we employed invertible Sentence-BERT networks [28] originally developed in cryptographic applications. The quality of the neural response-based predictions was evaluated using our ODIEU benchmark and was found to be significantly above chance. Together, our studies establish a computational pipeline that converts neural responses to odorants into language.

## Results

### Olfactory datasets

Our goal is to investigate the role of syntax in shaping human representations of odorants. Three of the four datasets discussed above include free-form descriptions of human olfactory percepts (Figure 1). These include Leffingwell [10], Arctander [12], and GoodScents [11] hereafter called LAG. Notably, LAG datasets overlap across a substantial number of odorants, meaning that many molecules are described by multiple independent observers (Figure 1).

Analysis of these overlaps reveals that 1,762/854 molecules have independent free-form descriptions in two/three datasets respectively, yielding 4,324 pairwise comparisons. For these molecules, discrete semantic labels-extracted by trained annotators to capture salient and widely used odorant features are also available. This data suggests that current datasets may contain sufficient information to quantify human odor perception.

### Dataset augmentation

Because the original human descriptions (HD) in the LAG data are typically object phrases rather than complete sentences, we used a large language model (Llama 3.3 70B-Instruct [32]) to convert them into completed human descriptions (CHD). The model was prompted as shown in Fig. 2A.

**Figure 2:**
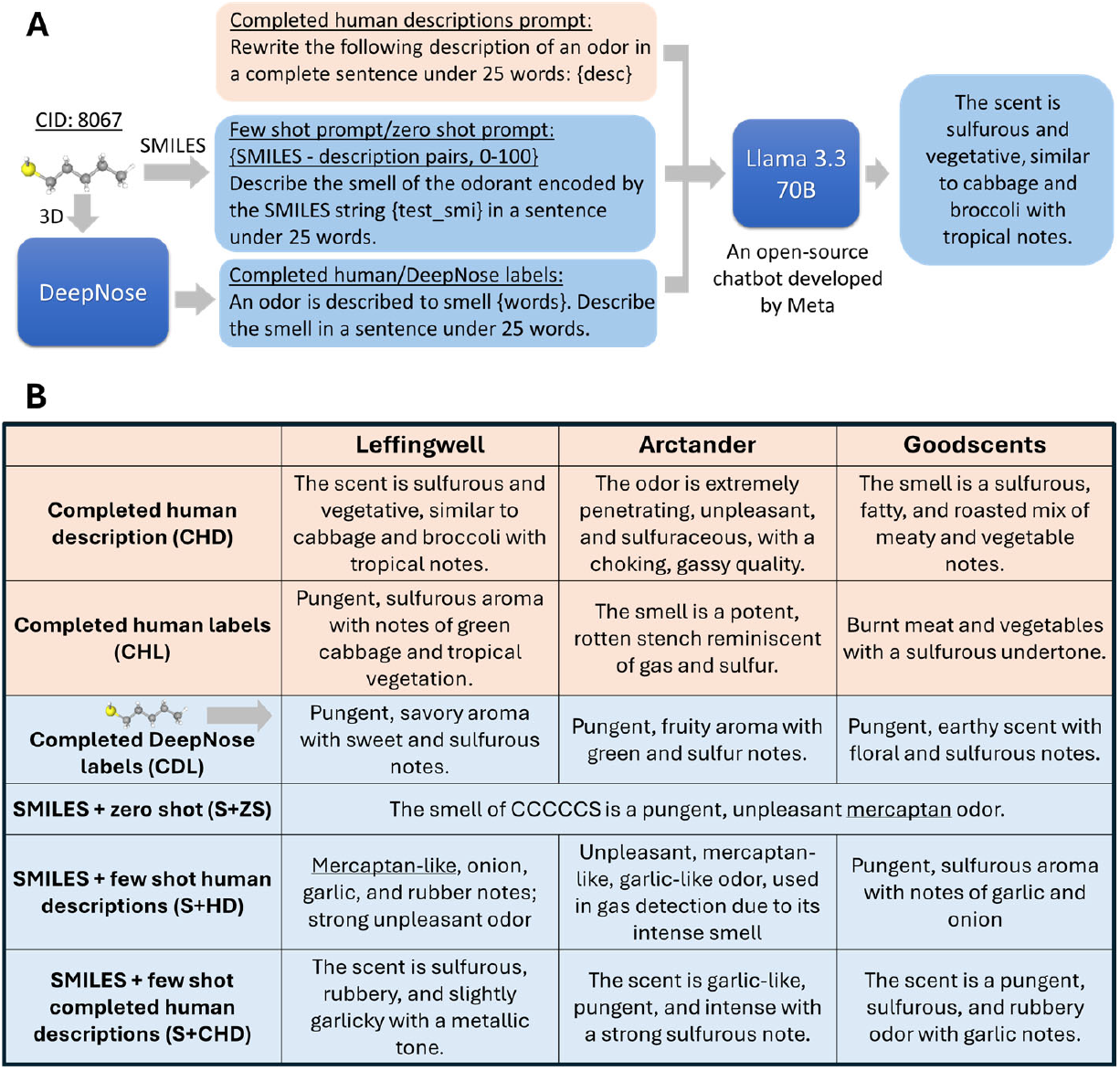
Dataset augmentation. (A) Because human descriptions (Figure 1) contained incomplete sentences, in many cases, missing both subject and verb, we augmented the dataset by generating short complete sentences (CHD) using Llama 3.3 [29]. To assess the importance of syntax in olfactory descriptions, we used Llama to generate completed human descriptions using human labels (CHL). In addition, we generated *de novo* descriptions using molecular structure alone. First, we asked Llama to generate descriptions using SMILES strings, either zero-shot or by providing several examples of SMILES-to-description matches in the prompt (few-shot). Second, we used the equivariant CNN developed earlier (DeepNose) to generate labels based on the 3D molecular structure. These labels were fed into Llama to yield completed sentences which we called completed DeepNose labels (CDL). (B) Examples of augmented descriptions for the molecule in Figure 1. Orange and blue backgrounds denote completed human descriptions and machine-generated sentences (using molecular structure or SMILES), respectively.

Using our model-based analysis (see below), we found that the generated CHDs remained close to the original HDs in sentence embedding space. Applying the same strategy, we also transformed human-provided labels (HL) into complete sentences (CHL). Because CHLs were generated using semantic information alone, the comparison between CHD and CHL descriptions will allow us to test whether adding syntax improves the quality of odor representations. Examples of these descriptions are highlighted in Figs. 2 and S1 (orange fields).

We next explored the generation of *de novo* text descriptions of odorant percepts from molecular structure alone. First, we prompted Llama 3.3 with examples of SMILES strings paired with human descriptions (HD, CHD, or CHL) for several molecules from our dataset and tasked it with generating descriptions for novel molecules. SMILES strings are a compact, human-readable format that encode molecular structure through atomic connectivity, making them suitable for direct input to language models. We varied the number of examples prompted from 0 to 1000 but observed no performance gains beyond 50 examples. As a result, we retained only the zero-shot and 50-shot conditions, referred to as SMILES+zero-shot (S+ZS) and SMILES+few-shot (S+CHD, S+HD, etc., Fig. 2). In the zero-shot condition, Llama generated odor descriptions based solely on its pre-trained knowledge of molecular structures and odors, without using any examples from our dataset. Second, we used DeepNose, an equivariant CNN trained on 3D molecular structures, to generate semantic descriptors. DeepNose achieves strong predictive performance (up to AUC = 0.91, depending on the dataset) [16, 17]. These semantic labels were then prompted to Llama to produce completed text descriptions, which we refer to as completed DeepNose labels (CDL, Fig. 2). The resulting *de novo* descriptions (S+ZS, S+CHD, S+HD, S+CHL, CDL) serve as test cases for evaluating our benchmark’s performance on novel S2T predictions.

### ODIEU dataset

Our dataset consists of various types of descriptions of percepts evoked by monomolecular odorants as illustrated in Figs. 1 and 2. It includes both original labels and free-text descriptions from the LAG dataset, as well as the augmented descriptions described above. For each molecule, we also provide structural information, including 3D atomic coordinates, .mol files, and SMILES strings, to facilitate downstream analysis and model development.

### Evaluating human odor descriptions using model free metrics

Our goal is to develop a metric for evaluating machine-generated S2T odor predictions. To this end, we leveraged the fact that our dataset includes descriptions for 4,324 pairs of molecules provided independently by two different human observers. These descriptions differ, resulting in an average human-to-human similarity score, denoted as *S*_*hh*_, which is smaller than the maximum possible similarity score (e.g., 1.0 if using a BLEU metric). A high-quality predictor should produce descriptions that resemble human descriptions as closely as two humans resemble each other, i.e., the average machine-to-human similarity score *S*_*mh*_ should approach *S*_*hh*_. In practice, machine-generated descriptions tend to diverge more, such that *S*_*mh*_*<S*_*hh*_. The gap between *S*_*hh*_ and *S*_*mh*_ thus provides a quantitative measure of prediction quality and allows to compare different S2T algorithms.

We first applied this logic to model-free N-gram-based measures of sentence similarity (BLEU [33], ROUGE [34], METEOR [35], BERTScore [36], Table 1). These metrics were computed by comparing completed human descriptions (CHDs) to other descriptions in the ODIEU dataset. For instance, the average BLEU-1 score between CHDs and completed human labels (CHLs) was approximately 51 (Table 1). In contrast, the BLEU-1 score between CHDs provided for the same molecule by two different humans was *S*_*hh*_ = 59, suggesting that free-form text descriptions of odorants (CHD) carry additional information about odor percepts beyond labels. Similarly, when comparing machine-generated descriptions based on molecular structure (CDLs) to human CHDs, the BLEU-1 score was *S*_*mh*_ = 46 (Table 1), i.e. lower than the human-to-human score. These findings demonstrate that the ODIEU dataset can serve as a benchmark for evaluating machine-generated odor descriptions.

**Table 1:** Results for evaluation of cross-human performance (columns CHD through CHL) and human CHD versus machine-generated descriptions (CDL through S+CHL). The abbreviations are defined in Figs. 1 and 2. For cross-human performance, we computed the metrics for each annotator’s CHD versus other annotators. For the generated descriptions, we computed the metrics for predictions versus human ground truth (CHD) for each molecule. All values are multiplied by 100 for brevity. CS stands for cosine similarity shown in Fig. 3. SEM values are included in parenthesis.

| Metric | CHD | HD | HL | CHL | CDL | S+ZH | S+HD | S+CHD | S+CHL |
| --- | --- | --- | --- | --- | --- | --- | --- | --- | --- |
| BLEU-1 | 59 (0.06) | 20 (0.06) | 9 (0.02) | 51 (0.06) | 46 (0.06) | 42 (0.04) | 20 (0.09) | 48 (0.08) | 56 (0.06) |
| BLEU-4 | 17 (0.05) | 1 (0.02) | 0 (0.00) | 10 (0.06) | 8 (0.02) | 5 (0.03) | 1 (0.02) | 8 (0.04) | 13 (0.04) |
| Rouge-L | 47 (0.04) | 20 (0.04) | 16 (0.04) | 35 (0.07) | 32 (0.03) | 25 (0.03) | 15 (0.04) | 35 (0.07) | 42 (0.04) |
| METEOR | 49 (0.04) | 16 (0.04) | 4 (0.01) | 39 (0.07) | 35 (0.03) | 30 (0.04) | 13 (0.06) | 38 (0.06) | 46 (0.05) |
| BERTScore (F1) | 92 (0.07) | 86 (0.08) | 82 (0.05) | 91 (0.07) | 90 (0.07) | 89 (0.07) | 86 (0.08) | 90 (0.04) | 92 (0.04) |
| ALL-MiniLM (CS) | 58 (1) | 53 (1) | 48 (1) | 53 (1) | 41 (2) | 0.33 (1) | 29 (2) | 32 (2) | 35 (2) |
| STELLA-400M (CS) | 55 (2) | 53 (2) | 48 (1) | 49 (2) | 33 (2) | 24 (2) | 24 (2) | 24 (2) | 26 (2) |

### Evaluating human descriptions using sentence embedding models

In addition to the model-free similarity measures described above, we evaluated representations of olfactory descriptions using sentence embedding networks. Our motivation was to develop a model that captures the specific characteristics of olfactory language present in our datasets. These embeddings enable us to address two key questions using domain-specific evaluators. First, do syntactic structures in odor descriptions convey additional information beyond simple word content? Second, how well do machine-generated descriptions align with human-authored ones when assessed in this sentence space? As the first step, we used two pretrained sentence-BERT (SBERT) models to compute sentence embeddings, ALL-MiniLM-L12-V1 [30] and STELLA-400M-v5 [31] with 33.4M and 435M parameters respectively. We found that both networks had hard time distinguishing descriptions of the same odorant from two randomly selected odorants (Fig. 3). For example, the cosine similarities between embeddings of descriptions of the same molecule from different datasets had a distribution that was strongly overlapping with the embeddings of two random molecules [Fig. 3B, Earth Mover Distance (EMD) of 0.14]. We also found that top-k retrieval accuracy for these embeddings is fairly low (38% for k=20, Fig. 3C). The average cosine similarities for descriptions of the molecule provided by two different datasets were comparable to those of two randomly chosen molecules (Fig. 3D, E). This observation implies that SBERT models pretrained on generic corpus emphasize non-olfactory content of the sentences.

**Figure 3:**
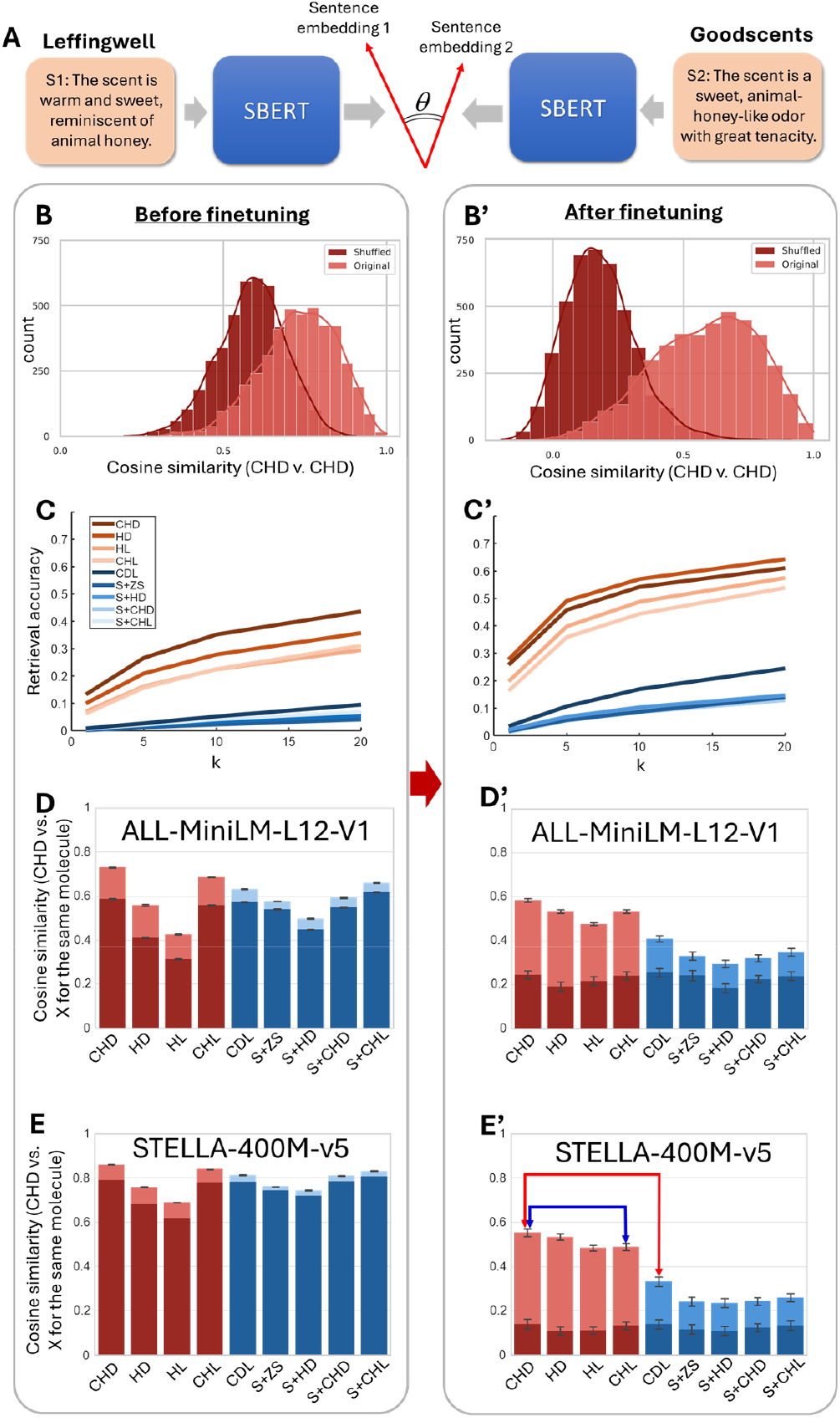
Model-based approach to evaluating the quality of olfactory descriptions. (A) Descriptions of the same molecule provided by different people (S1 and S2, c.f. Figure S1) were embedded into the sentence space using a sentence-BERT network (ALL-MiniLM-L12-V1 [30] or STELLA-400M-v5 [31]). The cosine similarity between sentence embedding was used to measure the diversity in human descriptions. (B-E’) Evaluating the quality of SBERT embeddings before (left) and after (right) fine-tuning with olfactory descriptions. (B) The distribution of CSs between descriptions of the same molecule by two different observers (lighter, CHD). The distribution of CSs has a strong overlap with CHDs of two different randomly chosen molecules [darker, shuffled, Earth Mover Distance (EMD) = 0.14], suggesting that pretrained SBERT models do not provide distinct embedding. (B’) After fine-tuning, the distribution of CS is more distinguishable from random (EMD = 0.40). (C) Retrieval accuracy rate for top-k ranked cosine similarities between SBERT embeddings of CHDs and SBERT embeddings of cross-dataset descriptions as indicated by color. The results are displayed for model STELLA-400M-v5 [31]. (C’) After fine-tuning, all retrieval accuracies are higher, indicating more distinguishable embeddings for each molecule. (D, E) Average CS between CHD embeddings of a molecule and the embeddings of the descriptions listed on the horizontal axis. Values for human-based descriptions (CHD through CHL) are shown by maroon bars, while S2T predictors are blue. Darker regions denote average CS for randomly shuffled molecules. Average CS between embeddings of the same molecule across datasets (lighter bars) are similar to randomly chosen molecules (darker bars). The pretrained models used are ALL-MiniLM-L12-V1 (D) and STELLA-400M-v5 (E). (D’, E’) After fine-tuning, average CSs are more distinguishable from random (light versus darker blue).

To teach SBERT to produce embeddings relevant to olfactory percepts, we fine-tuned the SBERT models using contrastive loss (AnglE loss [37]) using our dataset for training with 5-fold cross-validation. The two models displayed much better separation between pairs of descriptions from different datasets for the same versus different molecules (Fig. 3B’, EMD=0.40). Similarly, the top-k retrieval accuracy for fine-tuned embeddings was higher (Fig. 3C’, 64% for k=20). We found that the average CS for pairs of descriptions for the same molecule were dissimilar from random pairs (Fig. 3D’ and E’). Overall, we fine-tuned two SBERT models to produce text embeddings more distinguishable for different molecules in the olfactory context compared to the baseline models.

### Application of ODIEU benchmark: the role of syntax in odor description

Using our benchmark, we assessed the contribution of syntactic structure to odor prediction quality. Notably, the average CS between CHDs provided by different humans for the same molecule is higher than the average similarity between CHDs and CHLs (p<0.01, t-test, Fig. 3E’, blue arrow). Since CHLs are generated from human-provided semantic labels, this distinction indicates that syntactic structure encodes additional information relevant to odor perception. This finding underscores the need for the odor prediction models to move beyond semantic labels and better incorporate the syntactic richness of natural language descriptions.

### Using ODIEU to define the odor space

Perceptual odor spaces are actively investigated constructs in computational olfactory science [1, 3, 5-7, 16, 17, 19-24]. An odor space is defined as a representational space in which odorants are organized according to their perceptual effects on human subjects. Fine-tuned SBERT embeddings can be used to construct such an odor space. Because SBERT embeddings are highly multidimensional (D = 384 and 1024 for the ALL-MiniLM and STELLA models, respectively), we further reduced their dimensionality for visualization using the UMAP algorithm. We observed that odorants cluster according to perceptual families (Fig. 4A), suggesting that sentence embeddings may serve as a reasonable proxy for human olfactory perceptual space.

**Figure 4:**
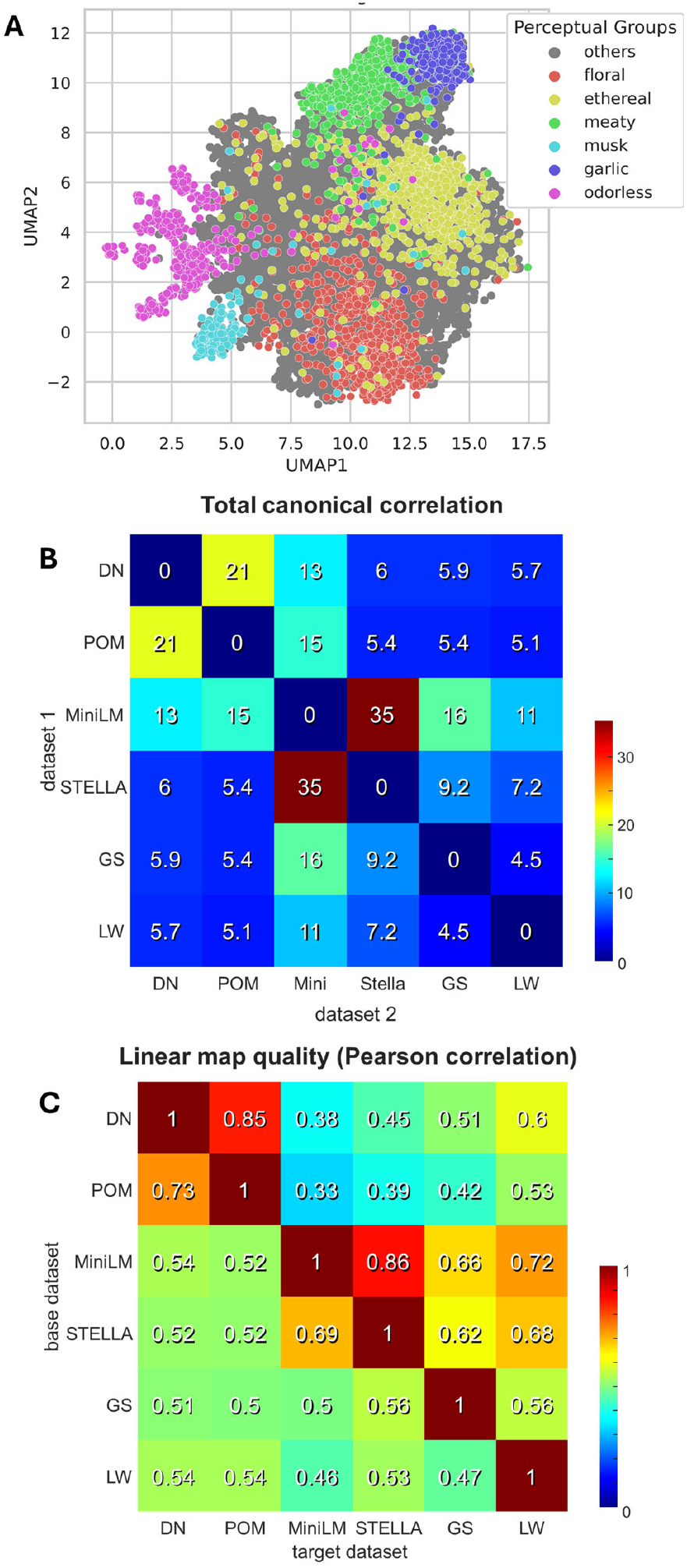
Odor space in the ODIEU embeddings. (A) SBERT odor space for the fine-tuned model (ALL-MiniLM-L12-V1). (B) Sum of squared canonical correlations for pairwise odor space comparisons between different spaces as indicated. (C) The results of linear regression from the base to target odor spaces quantified by the Pearson correlation over held-out data.

Because perceptual odor spaces have been defined multiple times in the past, we compared our SBERT embeddings with several existing models. Specifically, we examined the latent space of the DeepNose equivariant CNN [16, 17], the Principal Odor Map (POM) [9], and the Isomap embeddings of the GoodScents and Leffingwell datasets described previously [16]. Since the latent space of the POM model is not publicly available, we used OpenPOM instead, an open-source replica of the POM network available from Ref. [38] and displaying similar performance. Because the dimensionality of the SBERT embeddings is substantially higher than that of the other representations (384 and 1024 dimensions for MiniLM and STELLA, respectively), we projected them into 50-dimensional manifolds using the Multidimensional Scaling (MDS) algorithm [39], with minimal loss of information (MDS stress < 0.015). Overall, we analyzed six odor spaces: four generated by neural networks with different architectures (DeepNose, POM, MiniLM, and STELLA) and two derived from Isomap embeddings of human semantic datasets (GoodScents and Leffingwell).

To perform pairwise comparisons between odor spaces, we applied Canonical Correlation Analysis (CCA), which identifies pairs of linear projections of two datasets that maximize the total shared variance, defined as the sum of squared canonical correlations. This sum quantifies the effective number of shared dimensions between the two datasets, weighted by the strength of alignment in each dimension. In each case, the sum was computed over the set of overlapping molecules between each pair of odor spaces (Fig. 4B). For example, for the pair of DeepNose and POM embeddings, the total sum of squared canonical correlations was ∼21, indicating that these odor spaces share approximately 21 aligned dimensions. Similarly, for the representations generated by the fine-tuned MiniLM model compared with DeepNose and POM embeddings, the total sums of squared canonical correlations were 13 and 15, respectively. Overall, our results suggest that odor spaces derived using different algorithms and distinct types of data (semantic labels versus free-form text descriptions) exhibit substantial shared structure.

To further examine the relationships between different odor spaces, we asked whether odor coordinates in one space can be directly predicted from another. To address this question, we constructed predictive linear mappings between pairs of datasets (a base and a target space). Specifically, we estimated a linear transformation from the base dataset to the target dataset by computing the pseudoinverse transformation matrix using 80% of the molecules in the overlap. The remaining 20% of molecules were held out to evaluate prediction accuracy. Prediction performance was quantified as the Pearson correlation between the predicted odor coordinates and the true coordinates in the target space, computed on the held-out data. To prevent spurious correlations, each coordinate in both datasets was centered to zero using only the training set (80%) prior to fitting the mapping. Our results indicate substantial interdependencies across all datasets (Fig. 4). For example, predicting positions in the POM space from the DeepNose latent space yielded a correlation of R = 0.85. Similarly, MiniLM strongly predicted the STELLA space (R = 0.86). Notably, both fine-tuned SBERT embeddings were predictive of the GoodScents and Leffingwell spaces with high accuracy (R = 0.62-0.72). These latter spaces, Isomap embeddings derived from GoodScents and Leffingwell semantic labels, were not generated by neural networks and were also interrelated with both DeepNose and POM spaces. These findings suggest a mechanism by which deep learning networks (DeepNose and POM) generate perceptual predictions from molecular structure. The similarity between Isomap and network-based spaces indicates that neural networks partially recover the semantic structure of human odor perception captured by purely semantic embeddings, while also learning mappings from molecular structure into these perceptual spaces. Overall, our results demonstrate that independently derived odor spaces share substantial structure and can be predicted from one another with considerable accuracy.

### Evaluating simple S2T prediction models

We tested our model-based benchmark using two types of S2T predictions: SMILES-prompted and DeepNose label-based. As described in the Dataset Augmentation section, in the SMILES-based predictor, Llama 3.3 was prompted with a novel SMILES string to generate odor descriptions either without examples (S+ZS) or with 50 SMILES-to-CHD pairs (S+CHD) (Fig. 2B). S+ZS therefore reflects the competence of a general-purpose LLM in generating descriptions directly from SMILES strings, whereas S+CHD attempts to enhance this performance using in-context examples. Importantly, the prompts did not include the test molecule. To construct the DeepNose label-based descriptions (CDL), we first assigned semantic labels to odorants using our DeepNose network [17] and then prompted Llama 3.3 to generate complete text descriptions from these labels. Because the odorants for which the labels were generated belong to the DeepNose test sets, this procedure yields descriptions for previously unseen molecules.

We then applied our model-based metric to evaluate these S2T predictors. The average CS between embeddings of CHDs (ground truth) and CDLs (DeepNose-based predictions) was significantly lower than the average similarity between CHDs provided by different humans for the same molecule (Fig. 3E’, red arrow; also, Table 1). This indicates that DeepNose-based predictions do not match the quality of human descriptions. Similarly, SMILES-prompted descriptions (S+ZS, S+CHD, Fig. 3E’; Table 1) were of even lower quality, falling below both human and DeepNose-based outputs. These results demonstrate that our benchmark can effectively rate machine-generated odor predictions relative to human performance.

### Training an LLM to generate S2T odor descriptions

Our next goal was to design an algorithm capable of generating textual descriptions of odorant percepts directly from molecular structure. This task is analogous to image captioning or visual language grounding in computer vision. In the olfactory domain, it can be viewed as olfactory language grounding or S2T alignment. Given the strong performance of transformer-based autoregressive models, we adopted GPT-2 as our language backbone. However, because the number of available S2T training pairs is limited, a key challenge was to avoid degrading the pretrained language model during fine-tuning. To preserve its linguistic competence, we froze the GPT-2 weights and trained only a separate encoder. Specifically, we introduced a molecular encoder that takes representations of molecular structure and projects them into the token embedding space of GPT-2 (Fig. 5A). The resulting embedding was used as a learned prefix to condition the language model, enabling it to generate descriptions grounded in molecular structure. This prefix-tuning strategy is well suited for multimodal and conditional generation tasks, as prior work has demonstrated stable and reliable performance while preserving the integrity of the pretrained language model [40]. We call our algorithm and the resulting network Chemical Information Recognition and Annotation Network for Odors (CIRANO).

**Figure 5.**
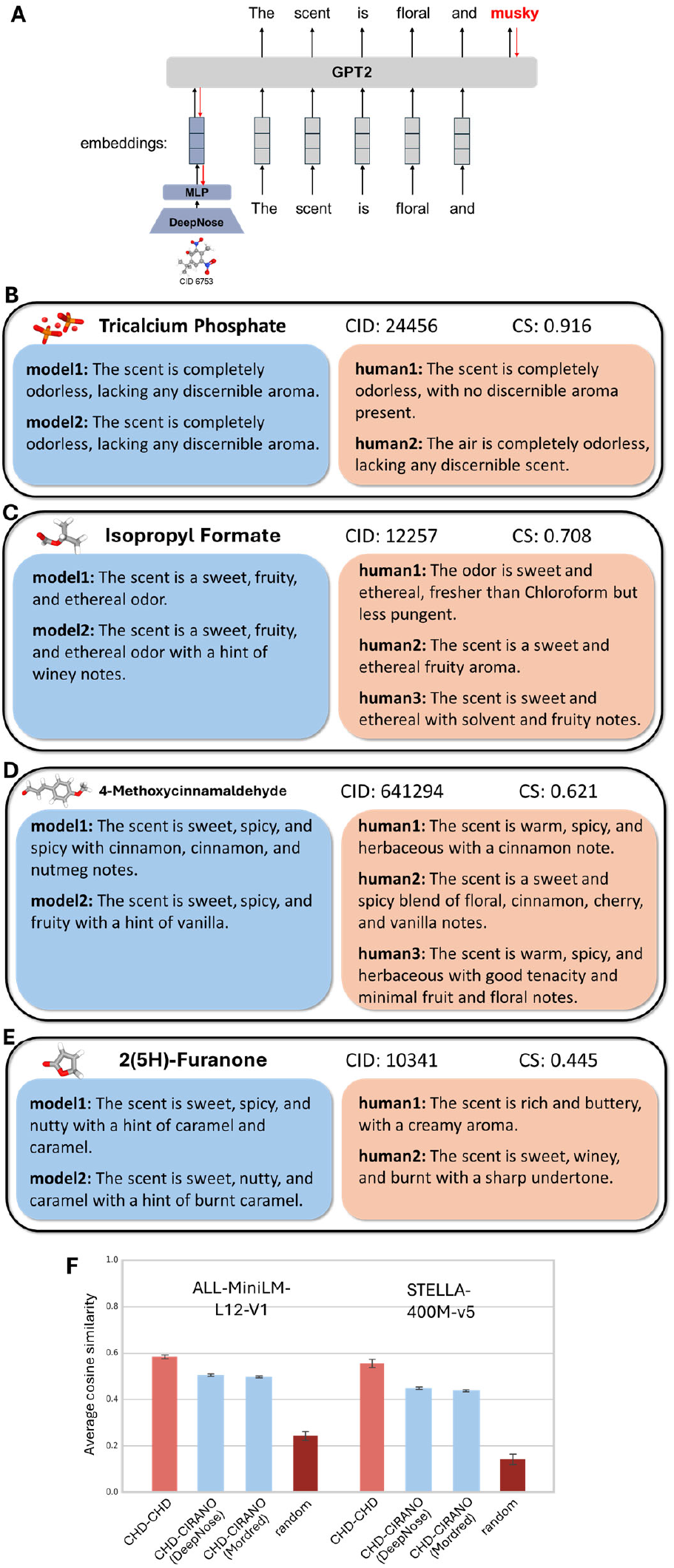
LLM that converts molecular structure into free-form textual odorant descriptions (CIRANO). (A) The structure of our network. (B-E) Examples of predictions generated by CIRANO from molecular structures. Model 1 and Model 2 were trained using DeepNose embedding vectors and Mordred fingerprints, respectively. (F) Application of our benchmark to CIRANO-generated predictions (blue), compared with human-human concordance (maroon).

The molecular structure in CIRANO was represented using two approaches. First, we used molecular shape representations extracted by the DeepNose network. These 96D vectors were generated based solely on molecular geometry, without explicitly incorporating bond information. Second, we used a set of 1,623 Mordred fingerprints for each molecule, *ad hoc* descriptors commonly used in computational chemistry to capture molecular shape and bond properties. These vectors were fed into a molecular encoder (a 4-layer MLP; see Methods), which projected them into the GPT-2 token space. During training, gradients were propagated through GPT-2 into the encoder network without updating GPT-2’s weights. This design mitigated the risks of overfitting and catastrophic forgetting when applied to conditional text generation from molecular representations.

The trained CIRANO model produces descriptions that are comparable to those of humans (Fig. 5B-E). Model fine-tuning largely preserved the natural language capabilities of the GPT-2 backbone, with rare exceptions (Fig. 5D), while enabling the generation of descriptions that capture a diverse set of olfactory percepts. We then applied the ODIEU benchmark to a held-out set of molecular descriptions by comparing CIRANO-to-human CS (CIRANO-CHD) with human-to-human one (CHD-CHD). For example, when the small MiniLM model was used in evaluation, CIRANO-CHD CS was 0.51 (0.003) with DeepNose embeddings used as inputs, compared to 0.58 (0.01) for CHD-CHD human concordance (Fig. 5F). Overall, although CIRANO predictions do not reach the level of human-human consensus, their scores were higher than those obtained using DeepNose labels alone (CDL). When Mordred fingerprints were used for prefix training, prediction quality was slightly lower [average CS = 0.499 (0.004)]. For the larger fine-tuned SBERT model used for evaluation (STELLA), CIRANO-CHD CS values were 0.45 (0.004) and 0.44 (0.004), compared to 0.55 (0.02) for CHD-CHD consensus. Overall, we trained CIRANO, an LLM capable of generating free-form natural-language descriptions of odorant percepts, that achieves performance comparable to, though slightly below, human-level consensus.

### Predicting odor descriptions using mouse neural data (N2T)

Our next goal was to develop an algorithm that predicts human odor text descriptions from neural responses. We call this type of algorithm a neural-to-text (N2T) prediction model. Because neural data for large odor sets is not available for humans, we used available mouse data [27], and called the descriptions generated from mouse neural data mouse-based descriptions (MD). The biological olfactory system relies on odorant receptors (ORs) to detect and identify airborne molecules. ORs are specialized proteins expressed by olfactory sensory neurons (OSNs) in the olfactory epithelium. Each OSN expresses a single OR type from a repertoire of hundreds in most vertebrates. OSNs expressing the same OR type project to discrete structures on the surface of the olfactory bulb (OB), called glomeruli, with each glomerulus receiving input from a single OR type. Odorants activate distinct patterns of glomeruli, determined by how different ORs respond to the molecule [41, 42]. These spatiotemporal activation patterns allow the olfactory system to encode odor identity. In our previous publication [27], we recorded responses from large glomerular arrays on the surface of the mouse OB to large odor sets in two individual mice. We embedded the glomerular odor responses into a relatively low-dimensional space (D = 30) without substantial loss of information. We showed that neural response embeddings in this space, called Q-space, generalize well across individual mice. Overall, the neural responses were represented in 30-dimensional Q-space, and the ODIEU dataset shared an overlap of N = 59 odors with the available mouse neural data. These embeddings served as input to our N2T model.

Training a CIRANO-like language model on mouse data was not feasible due to the small number of molecules (N = 59) for which mouse neural responses are available. Instead, we focused on building a predictive map between the mouse neural space and the SBERT embedding space. One option was to construct a mapping between the neural space and the fine-tuned SBERT embedding space. Indeed, our fine-tuned SBERT models generated high-dimensional vector embeddings for free-form human olfactory descriptions, and a predictive map between the mouse neural space and these embeddings could help link mouse and human responses to odorants. Fine-tuned SBERT models, however, provide only one-way text-to-embedding mapping and cannot be used to generate text descriptions. This limitation becomes problematic when predicting human-like text descriptions from mouse neural responses. To build an N2T model, we therefore relied on recently developed invertible SBERT models (invSBERT) [28, 43], which implement both text-to-embedding and embedding-to-text mappings and can be used as cryptographic encoder-decoder pairs.

To implement the N2T model, we constructed a linear transformation from the space of mouse neural responses, described by Q-vectors [27], to the invSBERT embedding space (Fig. 6A) using the Procrustes algorithm [44]. To validate our predictions, we used leave-one-out cross-validation (LOOCV), in which the Procrustes linear map is computed using all odorants except one and then applied to the left-out odorant. This procedure is repeated by leaving out each odorant in turn. As a result, we obtained and tested a predictive linear map between the space of mouse neural responses and the human invertible sentence space. The accuracy of this map was quantified by computing CS between embeddings of real human sentences and embeddings predicted from mouse neural responses for the same odorants. We found that the Procrustes algorithm aligns mouse neural data with natural language descriptions significantly better than in the case of randomly shuffled odorants (Fig. 6B).

**Figure 6.**
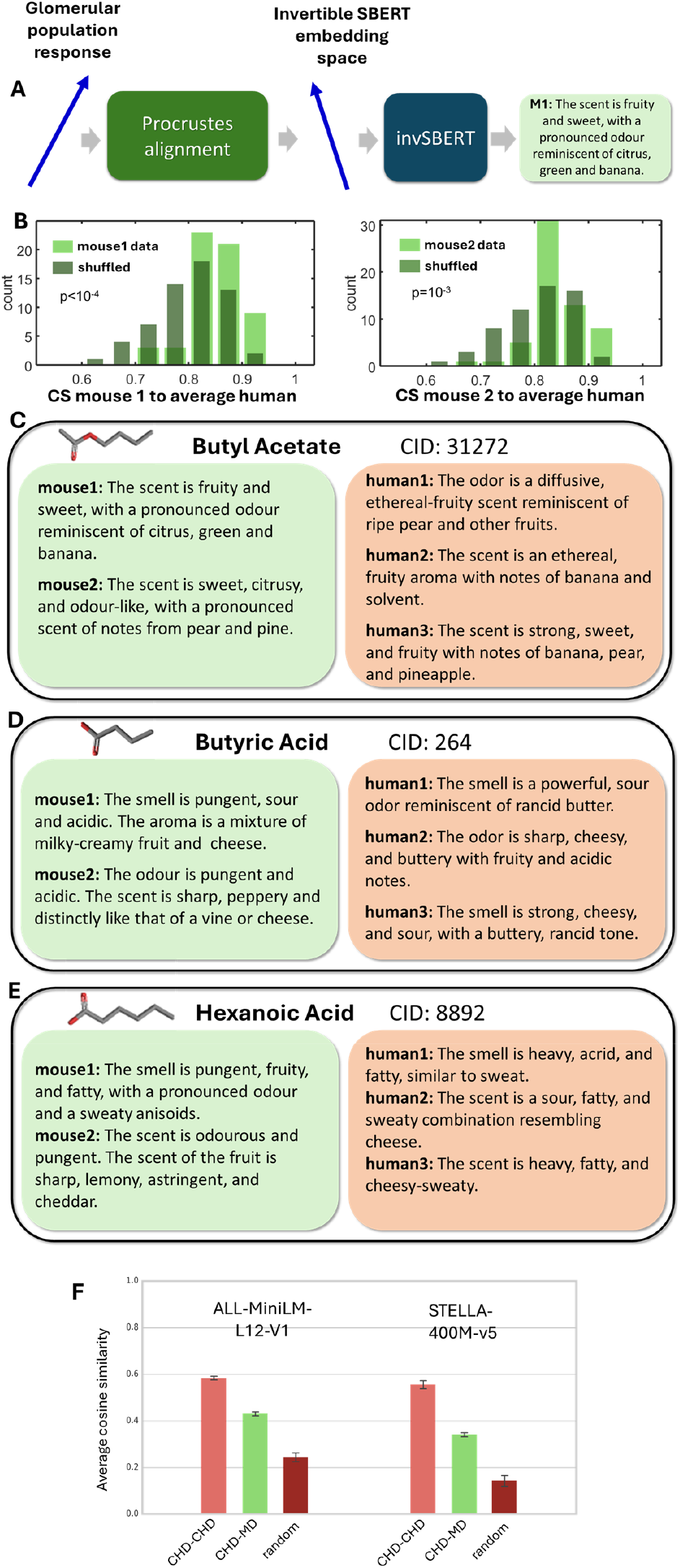
Neural-to-text (N2T) perceptual predictions. (A) The schematic for our computational pipeline. Glomerular population response vectors (Q-vectors [27]) were aligned with the invSBERT space using the Procrustes transform. The sentence embeddings produced based on neural responses were then inverted using the invSBERT model. (B) CS between invSBERT embeddings produced from neural responses and averaged over datasets CHD embeddings for the same odorant. Data for two mice is shown (left and right). Average CS is higher than for two random smells (t-test, p<10^-3^). (C-E) Examples of sentences generated from mouse data compared to original human CHDs for three molecules. (F) The CS of N2T predictor (CHD-MD, green) is lower than concordance of descriptions of two humans for the same molecule (light maroon, CHD-CHD) but higher than that for random human-human comparison (dark maroon).

Using invSBERT [45], we tested whether embeddings derived from neural data correspond to realistic human odor descriptions. A different pipeline with similar objective was recently introduced to map human neural responses to language [46]. Our results indicate that mouse neural data can be used to generate odorant text descriptions resembling those produced by humans (Fig. 6C-E). We applied our model-based benchmark to evaluate the quality of odor descriptions generated from mouse neural responses. We find that the average CS between mouse-derived and completed human descriptions (CHD) was 0.43 and 0.36 for the small and large fine-tuned SBERT models (MiniLM and STELLA), respectively. This implies that the mouse-to-human CS is lower than that for direct human-to-human comparisons (CS = 0.58 and 0.55; Fig. 6F, left bars), but significantly higher than chance (CS = 0.25 and 0.14; Fig. 6F, right bars). Overall, these findings indicate that mouse neural responses can be used to generate natural-language odor descriptions, although the quality of these texts does not yet match that of human generated descriptions.

### One person against the average

Several previous studies have shown that average panel responses are better predictors of individual percepts than another individual human. First, in profiling studies, it was noticed that averaged perceptual vectors become more stable when the number of individuals exceeds ∼50, suggesting that individual responses exhibit substantial variability [47]. Second, the application of the POM model yields performance that is better than the median individual human performance [6]. To address whether the average panelist is a better predictor of individual odor percepts than another individual, we computed the CS between the dataset-averaged CHD embedding and an individual embedding. We found that this CS is consistently higher than the individual-to-individual cross-dataset CS. For example, using the ALL-MiniLM model, the average-to-individual CS is 0.87 (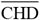, Table 2), which is substantially higher than the individual-to-individual CS of 0.58. Similarly, for the fine-tuned STELLA-400M model, the average-to-individual CS of 0.86 is higher than the cross-individual value of 0.55. This finding is not entirely surprising, however, because the dataset average includes the individual to which it is being compared. With only three datasets contributing to the average, this factor may artificially inflate the correlation. To address this issue, we computed an alternative average embedding that excludes the individual dataset being compared. For example, for CHDs drawn from the GoodScents dataset, we averaged CHD embeddings from the complementary Leffingwell and Arctander datasets. We denote these embeddings as 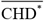(Table 2). The CS values computed using this strategy are lower than those obtained with the full average and are 0.61 and 0.58 for the fine-tuned MiniLM and STELLA models, respectively. These values are nevertheless slightly higher than the individual-to-individual CSs, suggesting that averaging across two individuals may yield a somewhat more accurate description embedding.

**Table 2:**
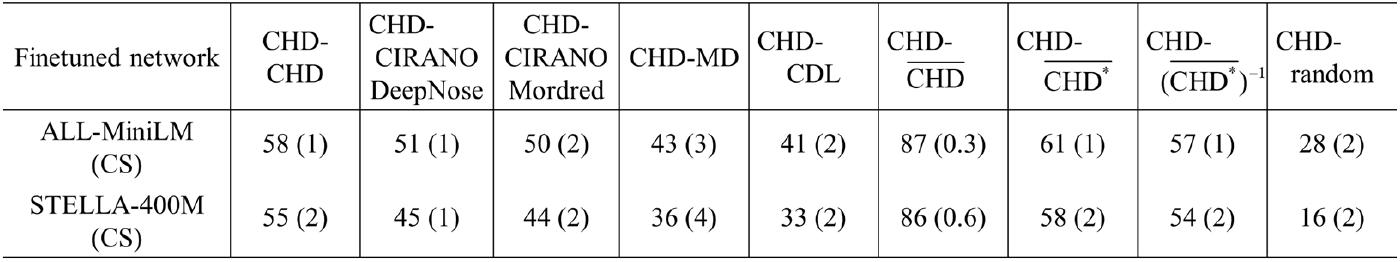
Results of S2T and N2T prediction models evaluated with the ODIEU model-based benchmark. The first column is also presented in Table 1. Results are also shown in Figs. 5 and 6. Notations are defined in the text. All values are multiplied by 100 for brevity. SEM values are included in parentheses.

This observation has several implications. First, it is possible that an S2T model may exceed the consensus between individual humans that we use as a benchmark. This may occur if the S2T model is trained to predict average responses. Second, this finding may indicate that odor descriptions are not perfectly accurate representations of perception due to individual cultural and genetic variability. Such variability is reduced when individual embeddings are averaged. We therefore hypothesize that an embedding averaged over many individuals may better capture the internal perceptual representation of individual humans.

Finally, we noted that these average embeddings may not correspond to a valid sentence. Because our benchmark is designed to evaluate sentences, we asked whether an average embedding can be used to generate a better odor description.

To this end, we computed the dataset-averaged invertible embeddings for each molecule and converted them into sentences using invSBERT. We then evaluated these sentences using our benchmark. As before, we excluded the dataset being tested from the average to avoid artificial inflation of the CS [Table 2, 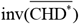]. We found that the descriptions generated from the average embeddings are of similar quality to the cross-individual ones (Table 2). This suggests that, although average embeddings may better predict individual human responses, sentences generated from such embeddings are not substantially better. Our benchmark therefore appears to provide a reasonable evaluation framework for models that generate natural language odor descriptions.

## Discussion

In this study, we assembled a dataset containing free-form text descriptions of odorant percepts and used this dataset to train language networks to generate odor descriptions. Our dataset consists of monomolecular odorants with a known 3D structure. We identified thousands of molecules for which descriptions are provided by more than one human observer. This data presented a unique opportunity to develop a benchmark for molecular S2T machine learning models. If the difference between machine-generated and human descriptions approaches the variability observed among human descriptions of the same molecule, one can argue that the Turing test for odorant S2T prediction is effectively met.

Based on this assumption, we developed our ODIEU benchmark through a three-step process. First, we assembled data, including augmented human descriptions of odorant percepts for thousands of molecules, into the ODIEU dataset. Second, we applied model-free N-gram-based sentence similarity metrics, such as BLEU, ROUGE, and METEOR, to quantify variability among human-generated descriptions. Finally, we fine-tuned transformer-based sentence embedding models of varying sizes to discriminate between random and human ground truth descriptions of molecules. The resulting average human-to-human similarity scores provide a quantitative target for evaluating S2T prediction algorithms.

Using the ODIEU benchmark, we made several observations. First, the differences between augmented human descriptions (CHD) and augmented semantic labels (CHL) for the same molecules are substantially greater than the differences between text descriptions provided by different human observers. This suggests that free-form human descriptions encode additional information about odorant percepts through their syntactic structure. Second, we found that a high-fidelity neural network predictor with strong classification performance (DeepNose, AUC ≈ 0.91) can generate realistic S2T descriptions. However, these predictions still fall short of matching the quality of human-generated descriptions. Together, these findings highlighted the limitations of current models and emphasized the need for benchmarks like ODIEU to guide the development of more linguistically and perceptually grounded odor prediction systems.

To overcome these limitations, we trained a specialized network, called CIRANO, to generate natural language odor descriptions directly from molecular structure. CIRANO uses a frozen GPT-2 backbone with a prefix network trained to incorporate olfactory domain knowledge. GPT-2 is a transformer-based autoregressive large language model introduced by OpenAI in 2019 [48] that contains 1.5 billion parameters and was trained on broad text corpora for next-token prediction and general language generation. By freezing its parameters, we leveraged its proficiency in English while embedding domain-specific information within the specialized prefix module. Similar algorithm has been found to be efficient in generating automatic image captions [40]. Using our benchmark, we found that CIRANO produces high-quality predictions with the greatest similarity to human descriptions among the S2T methods we tested. The highest accuracy was achieved when molecules were represented using DeepNose latent-space embeddings. Because these representations are assumed to approximate ensemble responses of human odorant receptors [16, 17], CIRANO may also provide insights into the putative mechanisms of olfactory processing. Overall, we trained an LLM to generate natural-language odor descriptions that approach the variability observed between human raters.

To explore alternatives to structure-based representations of odorants, we considered neural responses recorded in the olfactory system. Because large-scale neural recordings for diverse odor panels are not available in humans, we relied on mouse neural data. Given the relatively small number of odorants for which recordings were available (N = 59 in our dataset), we could not train a dedicated N2T network to generate human descriptions directly from these responses. Instead, we leveraged an invertible Sentence-BERT model [28], whose embedding space can be used to generate sentences. By mapping neural population activity into the invertible SBERT space, we generated human natural-language odor descriptions from mouse neural responses. The quality of the generated N2T descriptions did not reach human-human consistency or the performance of CIRANO, but it exceeded that of *ad hoc* baselines, such as SMILES- or DeepNose-based prompting methods. This level of performance is notable given the small number of molecules used to train the mapping. Expanding neural datasets and integrating them with ODIEU should further improve the quality of N2T predictions.

Our algorithms allow us to embed odorants into an odor space according to their effects on human observers. Research on odor spaces has spanned several decades and has successfully approximated olfactory perceptual spaces with manifolds of relatively low dimensionality. Recent deep learning models make it possible to map molecular structure onto perceptual spaces and to generate S2T predictions. In this study, we compared the subset of olfactory spaces and found that they share substantial similarities. For example, we showed that the previously published POM space can be predicted from DeepNose latent embeddings with high accuracy (R = 0.85, Fig. 4C). More generally, many odor spaces obtained using different methods shared a common structure (Fig. 4), as revealed by both CCA and predictive linear mappings. This finding suggests that diverse approaches to describing odorants converge on shared underlying dimensions. The relatively low dimensionality of olfactory manifolds suggests that olfaction may be analogous to vision: from the many physicochemical properties of odorants, the human olfactory system extracts only a small subset of the most relevant features. Deep learning models that generate S2T predictions, such as DeepNose, the POM model, or CIRANO, may help elucidate the significance of these dimensions.

In conclusion, we curated a large-scale dataset and developed ODIEU, a benchmark for evaluating the quality of natural-language odor descriptions. Using this benchmark, we showed that human odor descriptions encode perceptual information in their syntactic structure beyond discrete semantic labels. Leveraging this dataset, we trained a specialized large language model, CIRANO, to infer perceptual odor qualities directly from molecular structure and express them in natural language. Finally, we introduced a method for mapping mouse neural responses onto human odor descriptions. Together, CIRANO and ODIEU establish a framework for generating natural-language olfactory descriptions and quantitatively assessing their alignment with human perception.

## Acknowledgements

This work was supported by the National Institutes of Health BRAIN Initiative grant number U19NS112953, Swartz Foundation for Computational Neuroscience, and, in part, by National Science Foundation grant PHY-1748958 to the Kavli Institute for Theoretical Physics. We are thankful to Dmitry Rinberg and Hirofumi Nakayama for collecting and sharing mouse neural data.

## References

[1] A. Koulakov, B. E. Kolterman, A. Enikolopov, and D. Rinberg, “In search of the structure of human olfactory space,” (in English), Frontiers in Systems Neuroscience, Original Research vol. 5, 2011-September-15 2011, doi: 10.3389/fnsys.2011.00065.

[2] P. M. Wise, M. J. Olsson, and W. S. Cain, “Quantification of Odor Quality,” Chemical Senses, vol. 25, no. 4, pp. 429–443, 2000, doi: 10.1093/chemse/25.4.429.

[3] R. M. Khan et al., “Predicting Odor Pleasantness from Odorant Structure: Pleasantness as a Reflection of the Physical World,” The Journal of Neuroscience, vol. 27, no. 37, pp. 10015–10023, 2007, doi: 10.1523/jneurosci.1158-07.2007.

[4] H. Nakayama, R. C. Gerkin, and D. Rinberg, “A behavioral paradigm for measuring perceptual distances in mice,” (in eng), Cell Rep Methods, vol. 2, no. 6, p. 100233, Jun 20 2022, doi: 10.1016/j.crmeth.2022.100233.

[5] S. S. Schiffman, “Physicochemical Correlates of Olfactory Quality,” Science, vol. 185, no. 4146, pp. 112–117, 1974, doi: 10.1126/science.185.4146.112.

[6] B. K. Lee et al., “A principal odor map unifies diverse tasks in olfactory perception,” Science, vol. 381, no. 6661, pp. 999–1006, Sep 2023, doi: 10.1126/science.ade4401.

[7] M. Zarzo and D. T. Stanton, “Understanding the underlying dimensions in perfumers’ odor perception space as a basis for developing meaningful odor maps,” Attention, Perception, & Psychophysics, vol. 71, no. 2, pp. 225–247, 2009/02/01 2009, doi: 10.3758/APP.71.2.225.

[8] A. Dravnieks and A. Dravnieks, “Atlas of Odor Character Profiles,” in Atlas of Odor Character Profiles, vol. DS61–EB: ASTM International, 1992, p. 0.

[9] A. Keller et al., “Predicting human olfactory perception from chemical features of odor molecules,” (in eng), Science, vol. 355, no. 6327, pp. 820–826, Feb 24 2017, doi: 10.1126/science.aal2014.

[10] B. Sánchez-Lengeling, J. N. Wei, B. K. Lee, R. C. Gerkin, A. Aspuru-Guzik, and A. B. Wiltschko, “Machine Learning for Scent: Learning Generalizable Perceptual Representations of Small Molecules,” ArXiv, vol. abs/1910.10685, 2019.

[11] W. Luebke, “The Good Scents Company,” vol. http://www.thegoodscentscompany.com/. [Online]. Available: http://www.thegoodscentscompany.com.

[12] S. Arctander, Perfume and flavor chemicals (aroma chemicals). Montclair, N.J.,, 1969.

[13] T. Acree and H. Arn. “Flavornet and human odor space.” https://www.flavornet.org/ (accessed.

[14] S. Kim et al., “PubChem 2023 update,” Nucleic Acids Research, vol. 51, no. D1, pp. D1373–D1380, 2022, doi: 10.1093/nar/gkac956.

[15] E. A. Hamel et al., “Pyrfume: A window to the world’s olfactory data,” Scientific Data, vol. 11, no. 1, p. 1220, 2024/11/12 2024, doi: 10.1038/s41597-024-04051-z.

[16] N. Tran, D. Kepple, S. Shuvaev, and A. Koulakov, “DeepNose: Using artificial neural networks to represent the space of odorants,” presented at the Proceedings of the 36th International Conference on Machine Learning, Proceedings of Machine Learning Research, 2019. [Online]. Available: https://proceedings.mlr.press/v97/tran19b.html.

[17] S. Shuvaev, K. Tran, K. Samoilova, C. Mascart, and A. Koulakov, “DeepNose: An Equivariant Convolutional Neural Network Predictive Of Human Olfactory Percepts,” p. arXiv:2412.08747 doi: 10.48550/arXiv.2412.08747.

[18] F. Taleb, M. Vasco, A. Ribeiro, H. nio, M. a. r. Bjrkman, and D. Kragic, “Can Transformers Smell Like Humans?,” 2024. [Online]. Available: https://proceedings.neurips.cc/paper_files/paper/2024/file/848350afc07c02edea5be06d3390d865-Paper-Conference.pdf.

[19] B. Berglund, U. Berglund, T. Engen, and G. Ekman, “Multidimensional analysis of twenty-one odors,” Scandinavian Journal of Psychology, vol. 14, no. 1, pp. 131–137, 1973, doi: 10.1111/j.1467-9450.1973.tb00104.x.

[20] H. Chae, D. R. Kepple, W. G. Bast, V. N. Murthy, A. A. Koulakov, and D. F. Albeanu, “Mosaic representations of odors in the input and output layers of the mouse olfactory bulb,” Nature Neuroscience, vol. 22, no. 8, pp. 1306–1317, 2019/08/01 2019, doi: 10.1038/s41593-019-0442-z.

[21] H. Giaffar, S. Shuvaev, D. Rinberg, and A. A. Koulakov, “The primacy model and the structure of olfactory space,” PLOS Computational Biology, vol. 20, no. 9, p. e1012379, 2024, doi: 10.1371/journal.pcbi.1012379.

[22] J. Kowalewski, B. Huynh, and A. Ray, “A System-Wide Understanding of the Human Olfactory Percept Chemical Space,” Chemical Senses, vol. 46, 2021, doi: 10.1093/chemse/bjab007.

[23] A. Madany Mamlouk and T. Martinetz, “On the dimensions of the olfactory perception space,” Neurocomputing, vol. 58-60, pp. 1019–1025, 2004/06/01/ 2004, doi: 10.1016/j.neucom.2004.01.161.

[24] Y. Zhou, B. H. Smith, and T. O. Sharpee, “Hyperbolic geometry of the olfactory space,” Science Advances, vol. 4, no. 8, p. eaaq1458, 2018, doi: 10.1126/sciadv.aaq1458.

[25] W. W. Qian et al., “Metabolic activity organizes olfactory representations,” eLife, vol. 12, p. e82502, 2023/05/02 2023, doi: 10.7554/eLife.82502.

[26] B. Sanchez-Lengeling, Wei, J. N., Lee, B. K., Gerkin, R. C., Aspuru-Guzik, A., & Wiltschko, A. B. Leffingwell Odor Dataset (1.0) Zenodo. [Online]. Available: 10.5281/zenodo.4085098

[27] K. Samoilova, J. S. Harvey, H. Nakayama, D. Rinberg, and A. Koulakov, “Order code in the olfactory system,” bioRxiv, p. 2025.11.16.688691, 2025, doi: 10.1101/2025.11.16.688691.

[28] J. X. Morris, V. Kuleshov, V. Shmatikov, and A. M. Rush, “Text Embeddings Reveal (Almost) As Much As Text,” p. arXiv:2310.06816 doi: 10.48550/arXiv.2310.06816.

[29] A. Grattafiori et al., “The Llama 3 Herd of Models,” p. arXiv:2407.21783 doi: 10.48550/arXiv.2407.21783.

[30] H. Face. https://huggingface.co/sentence-transformers/all-MiniLM-L12-v1 (accessed.

[31] D. Zhang, J. Li, Z. Zeng, and F. Wang, “Jasper and Stella: distillation of SOTA embedding models,” p. arXiv:2412.19048 doi: 10.48550/arXiv.2412.19048.

[32] LLaMA 3.3 70B-Instruct. (2024). Meta Platforms, Inc., Menlo Park, CA. [Online]. Available: https://ai.meta.com/llama/

[33] K. Papineni, S. Roukos, T. Ward, and W.-J. Zhu, “BLEU: a method for automatic evaluation of machine translation,” presented at the Proceedings of the 40th Annual Meeting on Association for Computational Linguistics, Philadelphia, Pennsylvania, 2002. [Online]. Available: 10.3115/1073083.1073135.

[34] C.-Y. Lin, “ROUGE: A Package for Automatic Evaluation of Summaries,” Barcelona, Spain, July 2004: Association for Computational Linguistics, in Text Summarization Branches Out, pp. 74–81. [Online]. Available: https://aclanthology.org/W04-1013/. [Online]. Available: https://aclanthology.org/W04-1013/

[35] S. Banerjee and A. Lavie, “METEOR: An Automatic Metric for MT Evaluation with Improved Correlation with Human Judgments,” Ann Arbor, Michigan, June 2005: Association for Computational Linguistics, in Proceedings of the ACL Workshop on Intrinsic and Extrinsic Evaluation Measures for Machine Translation and/or Summarization, pp. 65–72. [Online]. Available: https://aclanthology.org/W05-0909/. [Online]. Available: https://aclanthology.org/W05-0909/

[36] T. Zhang, V. Kishore, F. Wu, K. Q. Weinberger, and Y. Artzi, “BERTScore: Evaluating Text Generation with BERT,” p. arXiv:1904.09675 doi: 10.48550/arXiv.1904.09675.

[37] X. Li and J. Li, “AnglE-optimized Text Embeddings,” p. arXiv:2309.12871 doi: 10.48550/arXiv.2309.12871.

[38] OpenPOM - Open Principal Odor Map. (2023). [Online]. Available: https://github.com/BioMachineLearning/openpom

[39] I. Borg and P. J. F. Groenen, Modern multidimensional scaling : theory and applications, 2nd ed. (Springer series in statistics). New York: Springer, 2005, pp. xxi, 614 p.

[40] M. Tsimpoukelli, J. Menick, S. Cabi, S. M. A. Eslami, O. Vinyals, and F. Hill, “Multimodal Few-Shot Learning with Frozen Language Models,” p. arXiv:2106.13884 doi: 10.48550/arXiv.2106.13884.

[41] R. W. Friedrich and S. I. Korsching, “Combinatorial and Chemotopic Odorant Coding in the Zebrafish Olfactory Bulb Visualized by Optical Imaging,” Neuron, vol. 18, no. 5, pp. 737–752, 1997, doi: 10.1016/S0896-6273(00)80314-1.

[42] B. Malnic, J. Hirono, T. Sato, and L. B. Buck, “Combinatorial Receptor Codes for Odors,” Cell, vol. 96, no. 5, pp. 713–723, 1999, doi: 10.1016/S0092-8674(00)80581-4.

[43] Y. Chen, H. Lent, and J. Bjerva, “Text Embedding Inversion Security for Multilingual Language Models,” p. arXiv:2401.12192 doi: 10.48550/arXiv.2401.12192.

[44] J. C. Gower and G. B. Dijksterhuis, Procrustes problems (Oxford statistical science series, no. 30). Oxford ; New York: Oxford University Press, 2004, pp. xi, 233 p.

[45] M. John Xavier, K. Volodymyr, S. Vitaly, and M. R. Alexander, “Text Embeddings Reveal (Almost) As Much As Text,” presented at the The 2023 Conference on Empirical Methods in Natural Language Processing, 2023. [Online]. Available: https://openreview.net/forum?id=EDuKP7DqCk.

[46] J. Tang, A. LeBel, S. Jain, and A. G. Huth, “Semantic reconstruction of continuous language from non-invasive brain recordings,” Nature Neuroscience, vol. 26, no. 5, pp. 858–866, 2023/05/01 2023, doi: 10.1038/s41593-023-01304-9.

[47] A. Dravnieks, “Odor Quality: Semantically Generated Multidimensional Profiles Are Stable,” Science, vol. 218, no. 4574, pp. 799–801, 1982, doi: 10.1126/science.7134974.

[48] A. Radford, J. Wu, R. Child, D. Luan, D. Amodei, and I. Sutskever, “Language Models are Unsupervised Multitask Learners,” 2019.

