## Supplementary Material for "Grounding olfactory perception in language: Benchmarks and models for generating natural language odor descriptions"

#### A Dataset details

The dataset published as part of the ODIEU benchmark is composed of 10,124 odorant molecules and associated olfactory descriptions and descriptors. For each odorant molecule we provide the original human authored descriptions, the manually extracted labels, the completed descriptions and a dataset augmentation from an odor percepts predictor and SMILES-prompted machine-generated descriptions.

##### A.1 Human authored descriptions

**Data collection.** We collected olfactory descriptions and labels of odorant molecules from three different sources. Two datasets, Leffingwell and Arctander, are available from the open source Pyrfume data collection [3], available in csv format. The Leffingwell dataset is a curated collection of odorant molecules expertly labelled with one or more odorant labels from a fixed lexicon for research in olfaction. The Arctander dataset is also a collection of molecules and associated odorant descriptors, curated by a professional perfumer.

A third dataset that we collected comes from a publicly available collection of odor molecules with associated odor descriptions from the Goodscents Company (https://thegoodscentscompany.com). We automatically parsed the web pages to extract the olfactory descriptions of odorant molecules, as well as olfactory labels. The odor descriptions and labels are provided by the molecule manufacturers directly.

##### A.2 Dataset augmentation with Llama 3.3

We employed Meta's Llama-3.3-70B-Instruct model (accessed March 2025 with Cluster 2) via OpenAI's API client with a temperature setting of 0.8. The LLM was provided with the system prompt: *"You are a helpful assistant that generates descriptions of odors based on provided words"*. To augment the LAG dataset, descriptions of molecules were generated using prompts following the standard role-based format (system/user/assistant) from the OpenAI ChatML template.

**Generating completed human description (CHD).** The original LAG dataset contained descriptions that were incomplete, often missing subjects or verbs. To build a dataset with consistent structural quality, we used the LLM to rewrite raw descriptions into complete sentences (CHD) using the prompt as shown:

> **Prompt:** Rewrite the following description of an odor in a complete sentence under 25 words: <u>Sulfurous, green, vegetative, cabbage, broccoli like; tropical notes</u>
> **LLM Response:** The scent is sulfurous and vegetative, similar to cabbage and broccoli with tropical notes.

**Generating completed human labels (CHL).** In order to directly compare the representations of molecules from free-text descriptions (HD) and binary labels (HL), we also used the LLM to construct sentence descriptions from ground truth labels (CHL). The following is an example description generated from labels for 1-Pentanethiol:

**Prompt:** An odor is described to smell cabbage, green, sulfurous, tropical, vegetable. Describe the smell in a sentence under 25 words.
**LLM Response:** Pungent, sulfurous aroma with notes of green cabbage and tropical vegetation.

| | Leffingwell | Arctander | Goodscents |
|---|---|---|---|
| **Molecule** | 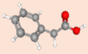 Phenylacetic Acid (CID: 999, SMILES: C1=CC=C(C=C1)CC(=O)O) | | |
| **Human description** | Sweet, animal-honey-like odor; honey, cocoa, floral taste | Sweet, animal-honey-like odor of extraordinary tenacity. The sweet-honey-like character is predominant at very low concentration, while the animal-Civet-like notes appear at higher concentrations. | sweet honey floral honeysuckle sour waxy civet. Luebke; Sweet, floral, honey, rose, chocolate, tobacco and powdery with animal nuances. Mosciano |
| **Human labels** | 'animal', 'honey', 'sweet' | 'animal', 'honey' | 'animal', 'chocolate', 'civet', 'floral', 'honey', 'honeysuckle', 'powdery', 'rose', 'sour', 'sweet', 'tobacco', 'waxy' |
| **Completed human description** | The scent is sweet, reminiscent of animal honey and floral notes. | The scent is a sweet, animal-honey-like odor with great tenacity. | The scent combines sweet honey and floral notes with a hint of sour and animal undertones. |
| 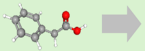 **Completed DeepNose labels** | Rich, sweet berry aroma with spicy and cheesy undertones, floral and green notes, on a woody base. | The scent is a vibrant, floral blend with fruity and spicy undertones. | Earthy, floral scent with sweet, woody, and fungal undertones. |
| **SMILES + zero shot** | The odor is acetic, pungent, and sour, like vinegar. | | |
| **SMILES + few shot completed human descriptions** | The scent features floral, herbal, and slightly sweet notes of violet. | The scent has a sweet, slightly fruity and herbal character. | The scent is a sweet, slightly spicy, and herbal phenolic aroma. |

**Figure S1.** Example of the dataset entry for a molecule (Phenylacetic Acid).

**Generating completed DeepNose labels (CDL).** To compare the performance of a classification model trained on olfactory datasets with the human-to-human benchmark, we generate *de novo* descriptions of molecules using labels from DeepNose, an equivariant CNN trained to map 3D molecular structures to semantic odor percepts [1]. Inputs to DeepNose consisted of 3D coordinates of C, H, O, N, S, Cl atoms for each molecule in the LAG dataset, obtained from PubChem based on their CID identifiers [2]. During inference, DeepNose generates outputs based on dataset-specific vocabularies due to a masking procedure applied during training (i.e., the same molecule may yield different predictions under the Leffingwell or Goodscents lexicons). Binary labels were assigned with threshold values that balance true and false positive rates computed on the training data.
The completed sentence descriptions were generated using the following template with DeepNose labels for 1-Pentanethiol using Leffingwell vocabulary as an example (underlined):

**Prompt:** An odor is described to smell fruity, meaty, onion, sulfurous, tropical, vegetable. Describe the smell in a sentence under 25 words.
**LLM Response:** Pungent, savory aroma with sweet and sulfurous notes.

**Generating SMILES-prompted zero-shot descriptions (S+ZS).** In order to assess generic LLM performance in olfactory description, we directly queried the model with SMILES strings to get zero-shot descriptions. The following is an example response for 1-Pentanethiol:

**Prompt:** Describe the smell of the odorant encoded by the SMILES string CCCCCS in a sentence under 25 words.
**LLM Response:** The smell of CCCCCS is a pungent, unpleasant mercaptan odor.

**Generating SMILES-prompted few-shot human descriptions (S+HD) and completed human descriptions (S+CHD).** In the few-shot setting, we provided the LLM with in-context example pairs of SMILES strings and ground truth sentences, using either raw human descriptions (HD) or completed descriptions (CHD) from the LAG dataset. To reduce potential bias in the description, examples were shuffled 50 times to generate descriptions for each SMILES string. After testing with varying numbers of examples provided for the task ranging from 1 to 500, we noticed no substantial improvement in cosine similarity to ground truth descriptions beyond approximately 50 examples. For the dataset and analyses, all few-shot descriptions used 50-shot responses.
The following two examples show LLM responses to few-shot prompting with either HD or CHD context, respectively.

**Examples:**
**Molecule:** CC=O
**Description:** Pungent ethereal-nauseating odor, in high dilution reminiscent of Coffee or Wine.
**Molecule:** CCOC(C)OCC
**Description:** Agreeable, refreshing, fruity-green odor.
**Molecule:** CC/C=C\CCOC(OCC/C=C\CC)C
**Description:** Powerful, oily-green odor with a sweet, vegetable-like undertone.
**Prompt:** Describe the smell of the odorant encoded by the SMILES string CCCCCS in a sentence under 25 words.
**LLM Response:** Mercaptan-like, onion, garlic, and rubber notes; strong unpleasant odor.

**Examples:**
**Molecule:** CC=O
**Description:** The scent is pungent and nauseating, similar to coffee or wine at high dilution.
**Molecule:** CCOC(C)OCC
**Description:** The scent is an agreeable, refreshing, and fruity-green odor.
**Molecule:** CC/C=C\CCOC(OCC/C=C\CC)C
**Description:** The scent is a powerful, oily-green odor with sweet undertones.
**Prompt:** Describe the smell of the odorant encoded by the SMILES string CCCCCS in a sentence under 25 words.
**LLM Response:** The scent is sulfurous, rubbery and slightly garlicky with a metallic tone.

#### A.3 Combined dataset

**Dataset availability.** The combined dataset, which we nicknamed LAG (for Leffingwell, Arctander, Goodscents), is available in a single file in csv format on the github repository (https://github.com/KoulakovLab/ODIEU/tree/main/datasets/descriptions.csv).

**Intended use.** The benchmark results are based on model training and comparison of generated descriptions over this dataset, using the completed human descriptions as reference. Future models for the prediction of semantic odorant percepts and natural language descriptions of odors may use this benchmark for training and compare their performance to other models and to natural language descriptions of odors authored by human beings. Since the completed human descriptions are used as reference, models should restrain from using this part of the dataset for training purpose or should take care of using the training/testing partition used for the finetuning of this benchmark's Sentence-

BERT models. This dataset can also be used to augment other olfactory datasets intended with the same aim.

**Training/testing set partitioning.** The models have been trained using the completed Leffingwell, Arctander and Goodscents descriptions, while the rest of the descriptions (human-authored, extracted labels and generated from DeepNose labels and SMILES prompts) have only been used for evaluating the performance of the Sentence-BERT models.

Furthermore, the completed human descriptions have been partitioned into five cross-validation folds. Each fold is composed of a training set (80% of available molecules) and a holdout, or testing, set (remaining 20%). The holdout sets have been chosen so that they tile perfectly the whole dataset without intersection, preventing contamination.

For the contrastive training of the Sentence-BERT models and the computation of cosine similarities, the pairs of descriptions have been chosen from the available molecules in the respective training and testing set. Since each training set contains descriptions for about 8,000 molecules, computing all possible pairs of descriptions was not a tractable operation. We therefore chose to include in the training set of each fold all the possible pairs of positive descriptions (descriptions of the same molecule from different sources, amounting for instance to 1883 pairs for the first fold) and randomly selected negative pairs of descriptions (descriptions of different molecules, amounting for instance to 37674 pairs for the first fold).

#### B Sentence-BERT Models

We publish two sets of Sentence-BERT models for all five folds that we specialized for separating olfactory descriptions of different molecules as part of the ODIEU benchmark. The models have been chosen based on their performance on standard benchmarks, as well as for their sizes. The number of parameters of the two models have been chosen to be one order of magnitude apart to show the effect of the size of the Sentence-BERT on the quality of the embeddings, on its performance pre- and post-finetuning.

##### B.2 Model parameters before finetuning

The pretrained (non-specialized on olfactory descriptions) version of the two models used in this benchmark are available on the Huggingface repository.

**Smaller model: all-MiniLM-L12-v1.** The pretrained model is available to download at https://huggingface.co/sentence-transformers/all-MiniLM-L12-v1. It contains 33.4 million parameters and has been trained on a variety of generic textual datasets (one billion sentence pairs) as part of a community event organized by Huggingface. This model maps sentences and paragraphs to a 384-dimensional dense vector space. The maximum token sequence length is 256; except for the original human-authored descriptions, all sequences are shorter than 100 tokens. More information on the details of the training of this model can be found on the model card following the link provided.

**Larger model: stella_en_400M_v5.** The pretrained model is available to download at https://huggingface.co/NovaSearch/stella_en_400M_v5. It contains 435 million parameters and has been trained on a variety of generic textual data. This model maps sentences and paragraphs to four dense vector spaces of dimensions 256, 512, 1024 and 12288 (the larger the embedding size the better the MTEB score). According to the authors, the difference in performance between the 1024 long embedding and the 12288 one is small (0.0001 on MTEB score); we therefore chose to work with the embedding of size 1024. The maximum token sequence length is 512 for this model; except for the original human-authored descriptions, all sequences are shorter than 100 tokens. More information on the details of the training of this model can be found on its Huggingface model card.

##### B.2 Finetuning procedure

The finetuned versions of these models can be downloaded as zip archives at https://github.com/KoulakovLab/ODIEU/tree/main/models/, and used for inference in python using the *sentence_transformers* library [4]. Each of the five models have been trained on a different cross-validation fold of our training set.

**Contrastive learning.** Pretrained models are downloaded and finetuned on olfactory data using the *sentence_transformers* python library. We chose to minimize the difference between a pair score (1 for positive pairs and 0 for negative pairs) and the AnglELoss between the embeddings of each description composing the pair. The two embeddings are obtained independently from the same Sentence-BERT model. The AnglELoss is computed using the *sentence_transformers.losses.AnglELoss* function from the *sentence_transformers* library, using a scale factor (hyperparameter) of 20.0.

**Training.** We trained each model for 50 epochs total, using a batch size of 128 per GPU and a warm-up ratio of 0.1 (ratio of total training steps used for a linear warmup from 0 to the learning rate). The optimizer is AdamW initialized with a learning rate of 5e-5. The final version of the model is chosen for the epoch in which the Kullback-Leibler divergence (KL-div) between the cosine similarities of the embedding of positive pairs and negative pairs is maximized on the validation set. Since the KL-div is non-symmetric, we considered three conditions: one where the reference distribution is the ordered similarities, one where the reference distribution is the shuffled cosine similarities and the average of the two. The first case showed the largest relative difference between ordered and shuffled computations of the cosine similarities and is the condition that we selected as stopping criterion for the training of our models.

**Cosine similarities of the embeddings.** The similarities between two embeddings e1 and e2 of olfactory descriptions are computed as

1.0 – *sklearn.metrics.pairwise.pairwise_distances*(e1, e2, metric="cosine")

using the pairwise_distances function from *scikit-learn* python library [5]. We computed the standard error of the mean by bootstrapping the mean cosine similarities obtained on each fold.

**Metrics.** We computed the benchmark metrics using the *evaluate* python library (https://pypi.org/project/evaluate/). We used the default parameters for each metric.

- BLEU: smoothing = False
- METEOR: alpha = 0.9, beta = 3, gamma = 0.5
- BERTScore: language = "en"

#### C Visualization of similarity scores for SBERT models

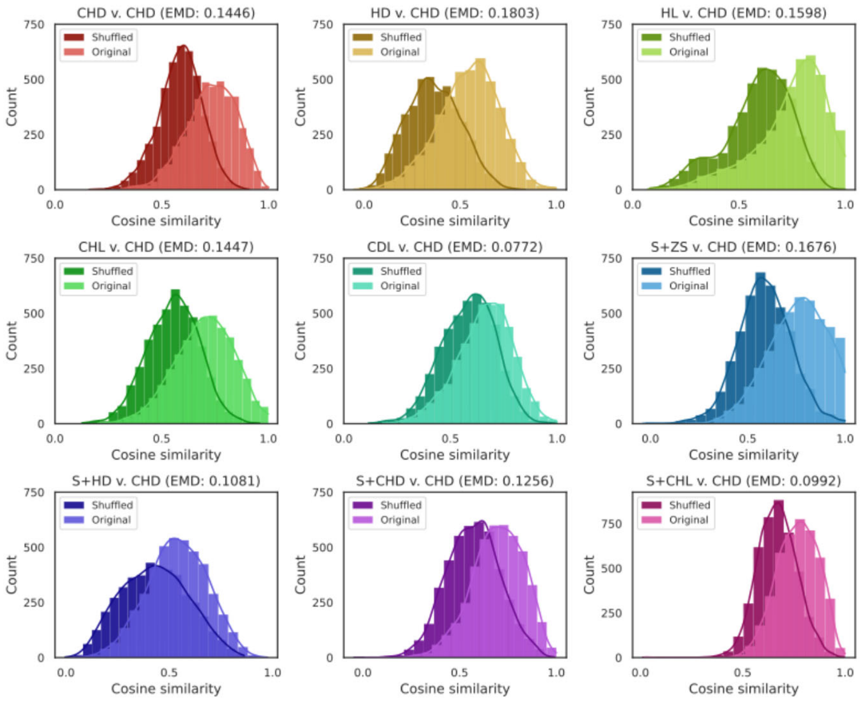


**Figure S2.** Cosine similarities between all types of descriptors and CHDs for base model with Earth Mover Distances (computed with *scipy.stats.wasserstein_distance*) between shuffled and original distributions.

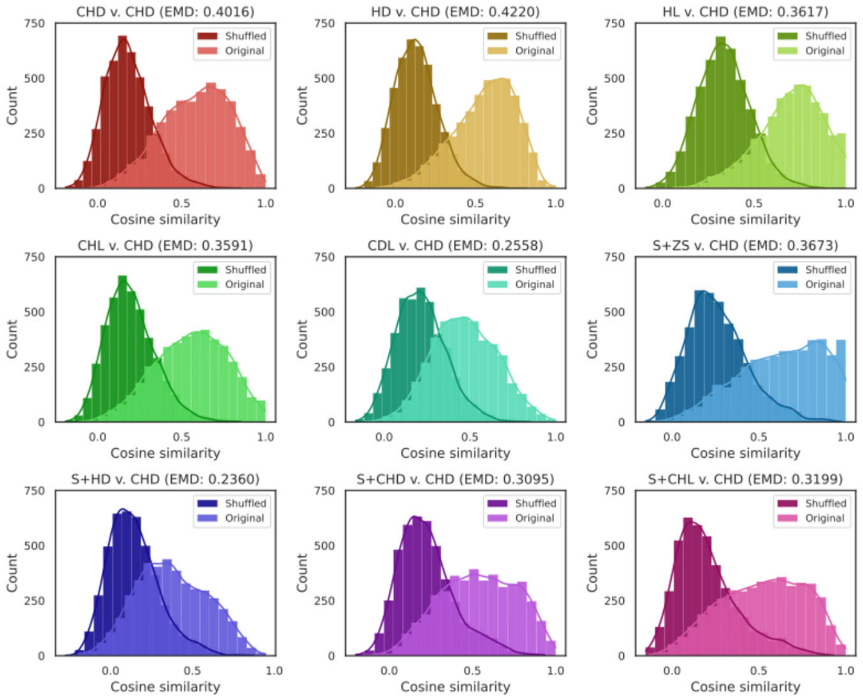


**Figure S3.** Cosine similarities between all types of descriptors and CHDs for finetuned model with Earth Mover Distances between shuffled and original distributions.

#### D UMAP visualization of SBERT embeddings

Sentence embeddings for CHDs were computed using the test molecules from each of the five cross-validation folds of which the SBERT models were trained on, resulting in a high-dimensional representation of the perceptual space. To visualize this embedding space, we applied UMAP as implemented in the *umap_learn* library with parameters `n_neighbors=100` and `min_dist=0.99` to reduce the vectors to two dimensions [6]. The resulting projections were labeled according to six perceptual odor groups. In particular, the group labels are "floral" (also includes descriptors such as "jasmine", "rose", "lavender"), "meaty" (e.g., "savory", "beefy", "roasted"), "ethereal" (e.g., "cognac", "fermented", "alcoholic"), "musk", "garlic", and "odorless". The results are displayed in Figure 4E.

### E CIRANO model description

We consider the conditional generation task of aligning molecular representation with natural language descriptions. The model that we focus on is based on freezing a pretrained language model and conditioning it via olfaction-specific continuous prefixes [1]. In this framework, a pretrained transformer decoder is conditioned on task-specific embeddings without changes to its internal model weights. Prefix encodings are produced by trainable external encoders that map from different molecular representations (DeepNose or Mordred) to fixed-length vectors that are prepended to the decoder input [2, 3].

The use of frozen pretrained language models is suitable for this task for several reasons. First, we rely on existing LLMs such as GPT2 that already encode robust semantic structure acquired from large scale text corpora [4]. Rather than attempting to further improve the performance of models on text generation, our objective is to preserve the pretrained model's linguistic competence and align its latent space to incorporate molecular information using efficient tuning strategies. Second, the conditioning of molecular bias via prefix tuning is an appropriate choice as prior works have noted stable and reliable performance for tasks in multimodal and conditional generation settings [1]. By training an encoder to produce a continuous prefix that conditions the frozen decoder, the model can integrate molecular information without perturbations to the internal linguistic representations of the pretrained language model. Finally, given the relatively limited size of available training data and the domain specificity of the task, this design choice mitigates the risks of overfitting and catastrophic forgetting when applied to the task of conditional text generation from molecular representations.

**CIRANO Architecture**

Our model, CIRANO, is comprised of a frozen pretrained GPT2 decoder and a trainable molecular encoder MLP. The molecular encoder maps a fixed-length molecular feature vector into a sequence of continuous embeddings of a specific length and a dimensionality which matches the hidden size of GPT2. The encoder uses a standard multilayer feedforward architecture parameterized by depth, hidden size, and dropout rate. The output prefix is also normalized using layer normalization and multiplied by a learned scaling factor. This scaling and normalization mechanism is critical for stabilizing prefix conditioning and is aligned with practices for conditioning autoregressive models in multimodal settings [1]. During training, the molecular prefix is concatenated with token embeddings from the GPT2 tokenizer to form a composite embedding sequence. The standard GPT2 tokenizer was used with the addition of a start token and adopting the end-of-sequence token for padding purposes. Then GPT2 decoder processes the unified sequence with an attention mask that does not distinguish between prefix and token positions. With the generated text, cross-entropy loss is computed with prefix positions masked to exclude them from gradient contributions.

**Dataset**

Testing and training experiments used the combined dataset, which for each molecule contains its corresponding molecular embeddings and sentence descriptions.

**Hyperparameter Optimization and Final Training**

For each choice of encoder, we implemented nested cross-validation to optimize parameters for the model. Using the same five data splits defined in the SBERT finetuning procedure as the outer fold, the remaining data is split into four inner folds. The inner folds of data were then used for hyperparameter optimization via the Optuna library with the objective of best average performance across splits [5]. The AdamW optimizer was used to train models for each trial. The most optimal parameters for each outer fold were selected as those that minimized the average loss on the validation sets across the inner folds.

To train the model and produce the final descriptions on the test set, the median was taken over the five set of best parameters for the outer folds. The final parameters used for each molecular embedding type are as followed:

| | DeepNose Encoder | Mordred Encoder |
|---|---|---|
| Input size | 96 | 1613 |
| Learning rate | 0.000368 | 0.000184 |
| Dropout | 0.275225 | 0.154602 |
| Number of MLP layers | 4 | 4 |
| MLP hidden size | 768 | 896 |
| Prefix length | 14 | 15 |
| Batch size | 8 | 8 |
| Epochs | 17 | 13 |

**Table S1:** Encoder parameters for CIRANO.

**Text Generation**

To generate a description for a molecule, a corresponding prefix is generated from the trained encoder using either DeepNose or Mordred features. The sequence is then initialized with the <START> token. We used a generic greedy decoding scheme where successive tokens were selected as the most probable token from a distribution over the vocabulary produced by the model. The sequence is terminated when either the end-of-sequence token is produced or the maximum number of new tokens has been reached (20 was the chosen value since this roughly corresponds to an average sentence length of 10-15 words for descriptions in the dataset).

**Implementation and Reproducibility**

All testing and final model training used PyTorch and the HuggingFace Transformers library. The dataset was split into 5 folds with consideration to avoid contaminating the same molecule appearing in both train and test set. Seeds were fixed for NumPy and PyTorch (CPU and GPU) to ensure reproducibility.

#### F Visualization of CIRANO predictive performance across percepts

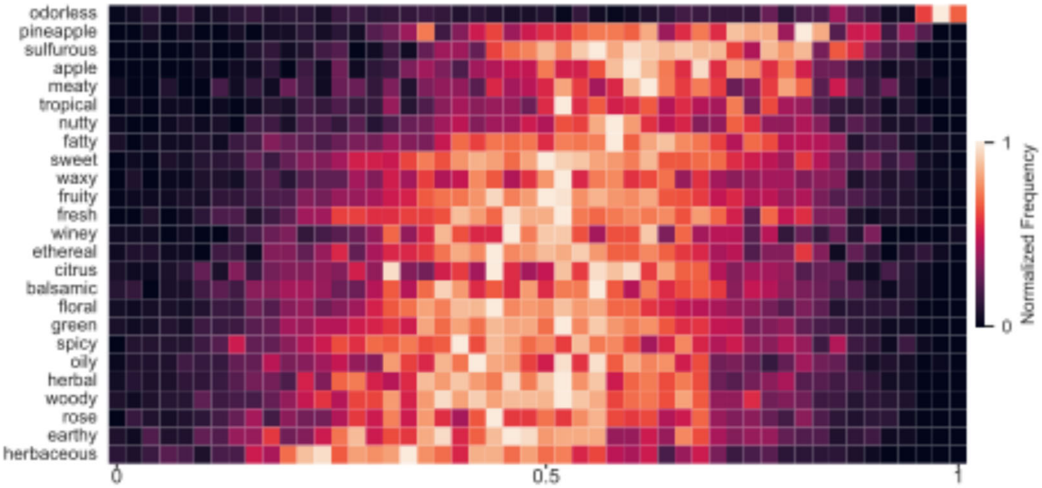


**Figure S4.** Heatmap showing the top 25 most frequently occurring labels (HL) in the dataset and CS values between generated descriptions and the corresponding ground truth text.

#### G Training resources

Training time, memory, CPU and GPU have been measured using the *psutil* and *gpustat* python libraries. Table S2 summarizes the resource usage for training and inference of the smaller and larger model.

The models have been trained using two GPU clusters. The smaller model (all-MiniLM-L12-v1) has been trained on cluster 1, the larger model (stella_en_400M_v5) on cluster 2. The hardware and software specifications of clusters 1 and 2 are summarized in Table S3. Total training time of over the five folds of the smaller model was 242 mn and 108 mn for the larger model. Three GPUs were available for the training of the smaller model, seven GPUs were available for the training of the larger model.

**Python libraries.** The complete list of necessary python libraries is available in the requirements file of the github repository https://github.com/KoulakovLab/ODIEU/blob/main/requirement.txt. We used the *sentence_transformers* library for downloading and training the models, as well as the accelerate library. We used the *scikit-learn* library for the computation of the cosine similarities. For the computation of the benchmark's metrics, we used the *evaluate* library from Huggingface.

| Model | CPU time (hours) | CPU utilization (%) | Memory (GiB) | GPU time (hours) | GPU utilization (%) |
|---|---|---|---|---|---|
| Smaller | 2.33 | 45 ± 21 | 5.6 ± 0.2 | 12.1 | 74.2 ± 34.8 |
| Larger | 2.4 | 45 ± 28 | 5.9 ± 0.4 | 12.6 | 49 ± 40 |

**Table S2:** Resource usage for the finetuning and embedding computation of the smaller model (all-MiniLM-L12-v1) and the larger model (stella_en_400M_v5).

| | **Cluster 1** | **Cluster 2** |
|---|---|---|
| **Operating system** | CentOS Linux 8 (Core) | Ubuntu 22.04 |

| | | |
|---|---|---|
| **Processors** | 80 × Intel® Xeon® Gold 6248 | 96 × AMD EPYC 7F72 24-Core |
| **Memory (RAM)** | 1535 GiB | 1,008 GiB |
| **Graphic processors** | 6 × Tesla V100S-PCIE-32GB | 8 × NVIDIA A100-SXM4-80GB |
| **Python version** | 3.11 | 3.12 |
| **Pytorch+CUDA version** | 2.7.0 + 12.6 | 2.6.0 + 12.4 |

**Table S3:** Hardware specifications of the two clusters used to train the models.

#### H Benchmark availability

The dataset, models and code are available on the https://github.com/KoulakovLab/ODIEU github repository.

##### References

[1] S. Shuvaev, K. Tran, K. Samoilova, C. Mascart, and A. Koulakov, "DeepNose: An Equivariant Convolutional Neural Network Predictive Of Human Olfactory Percepts," p. arXiv:2412.08747doi: 10.48550/arXiv.2412.08747.

[2] S. Kim *et al.*, "PubChem 2023 update," *Nucleic Acids Research,* vol. 51, no. D1, pp. D1373-D1380, 2022, doi: 10.1093/nar/gkac956.

[3] Jason B. Castro, *et al.*, “Pyrfume: A Window to the World’s Olfactory Data”, bioRxiv 2022.09.08.507170; doi: https://doi.org/10.1101/2022.09.08.507170

[4] Reimers, N., & Gurevych, I. (2019). “*Sentence-BERT: Sentence Embeddings using Siamese BERT-Networks"*. Conference on Empirical Methods in Natural Language Processing.

[5] Pedregosa, F., Varoquaux, Ga"el, Gramfort, A., Michel, V., Thirion, B., Grisel, O., … others. (2011). Scikit-learn: Machine learning in Python. *Journal of Machine Learning Research*, *12*(Oct), 2825–2830.

[6] L. McInnes, J. Healy, and J. Melville, "UMAP: Uniform Manifold Approximation and Projection for Dimension Reduction," arXiv:1802.03426, 2020, doi: 10.48550/arXiv.1802.03426.
